# Characteristics and functions of infection-enhancing antibodies to the N-terminal domain of SARS-CoV-2

**DOI:** 10.1101/2023.09.19.558444

**Authors:** Ruth I. Connor, Mrunal Sakharkar, C. Garrett Rappazzo, Chengzi I. Kaku, Nicholas C. Curtis, Seungmin Shin, Wendy F. Wieland-Alter, Joshua A. Weiner, Margaret E. Ackerman, Laura M. Walker, Jiwon Lee, Peter F. Wright

## Abstract

Characterization of functional antibody responses to the N-terminal domain (NTD) of the SARS-CoV-2 spike (S) protein has included identification of both potent neutralizing activity and putative enhancement of infection. Fcγ-receptor (FcγR)-independent enhancement of SARS-CoV-2 infection mediated by NTD-binding monoclonal antibodies (mAbs) has been observed *in vitro*, but the functional significance of these antibodies *in vivo* is not clear. Here we studied 1,213 S-binding mAbs derived from longitudinal sampling of B-cells collected from eight COVID-19 convalescent patients and identified 72 (5.9%) mAbs that enhanced infection in a VSV-SARS-CoV-2-S-Wuhan pseudovirus (PV) assay. The majority (68%) of these mAbs recognized the NTD, were identified in patients with mild and severe disease, and persisted for at least five months post-infection. Enhancement of PV infection by NTD-binding mAbs was not observed using intestinal (Caco-2) and respiratory (Calu-3) epithelial cells as infection targets and was diminished or lost against SARS-CoV-2 variants of concern (VOC). Proteomic deconvolution of the serum antibody repertoire from two of the convalescent subjects identified, for the first time, NTD-binding, infection-enhancing mAbs among the circulating immunoglobulins directly isolated from serum (*i.e*., functionally secreted antibody). Functional analysis of these mAbs demonstrated robust activation of FcγRIIIa associated with antibody binding to recombinant S proteins. Taken together, these findings suggest functionally active NTD-specific mAbs arise frequently during natural infection and can last as major serum clonotypes during convalescence. These antibodies display diverse attributes that include FcγR activation, and may be selected against by mutations in NTD associated with SARS-CoV-2 VOC.

## INTRODUCTION

Identification of functional antibody responses to the severe acute respiratory syndrome coronoavirus-2 (SARS-CoV-2) spike (S) protein has contributed to an understanding of the immune correlates of protection and informed vaccine development. The S protein of SARS-CoV-2 projects from the virion surface as a trimer of heterodimers, each consisting of two subunits, S1 and S2, formed by furin cleavage. Within the S1 subunit, the receptor-binding domain (RBD) mediates binding of the virus to angiotensin-converting enzyme-2 (ACE2) on host cells, and elicits neutralizing antibodies following natural infection and after vaccination (1–4). The RBD is subject to mutation contributing to the emergence of SARS-CoV-2 variants with increased transmissibility and significantly reduced susceptibility to neutralization (5–7). As the S protein constitutes the focus of many current vaccine platforms, considerable effort has been made toward understanding the evolution of the antibody repertoire and the functional impact of RBD mutations on neutralization of emerging SARS-CoV-2 variants of concern (VOC) (8).

Less well understood is the functional role and immunogenicity of the surface-facing N-terminal domain (NTD) of the S1 subunit. The NTD has multiple glycosylation sites among five highly variable loop structures (N1-5) that serve to fine tune and facilitate virus entry into host cells (9–12). NTD-binding antibodies can neutralize SARS-CoV-2 and these antibodies recognize epitopes within a convergent antigenic supersite (13–17), but also within conserved epitopes outside this site (18). Mutations in the NTD are present in all SARS-CoV-2 VOC to date and encompass residues constituting the neutralization supersite (14,15) and those predicted to contribute to evasion of serum neutralizing antibodies (17), consistent with immune selection driving evolution of the virus in this subdomain (8).

A small number of NTD-binding antibodies have been found to enhance SARS-COV-2 infection *in vitro* through an Fcγ-receptor (FcγR)-independent mechanism (19,20). These infection-enhancing antibodies recognize epitopes in the NTD outside the neutralization supersite and adjacent to the NTD variable loops (19,20). The significance of NTD-binding, infection-enhancing antibodies *in vivo* is less clear, as animal studies found minimal evidence of disease enhancement following antibody infusion and viral challenge in mouse and non-human primate models (20). These results are supported by evidence that NTD-binding antibodies can have additional functional attributes including FcγR-mediated effector functions that may contribute to viral clearance and protection from disease (21). However, detection of NTD-binding, infection-enhancing antibodies at a higher frequency in the sera of COVID-19 patients with severe disease has suggested a correlation between enhancing antibodies and disease severity (19). Overall, these findings leave unanswered questions as to the significance and functional attributes of infection-enhancing antibodies directed to the NTD.

In the present study, we sought to determine the frequency, characteristics and functional attributes of a panel NTD-binding antibodies derived from a large pool of S-specific mAbs from eight COVID-19 convalescent donors (22). Our findings demonstrate a broad antibody response to the NTD with evidence of neutralizing and infection-enhancing activities associated with mAbs from donors with mild or severe COVID-19. We present evidence of loss of antibody functionality to Omicron BA.1 and BA.2 variants that correlates with loss of binding to the variant S proteins. Moreover, we demonstrate for the first time the presence of NTD-binding, infection-enhancing antibodies as major clonotypes in the serum repertoire of two subjects, and show the corresponding recombinant mAbs have robust FcγR-activating functionality suggesting their potential to contribute to immune protection.

## RESULTS

### Frequency and binding specificity of SARS-CoV-2 infection-enhancing mAbs

The frequency of FcγR-independent, infection-enhancing antibodies to SARS-CoV-2 was determined by measuring the neutralizing or infection-enhancing activities of 1,213 mAbs derived from a previous longitudinal study of B-cell evolution in eight COVID-19 convalescent donors (22). Antibodies were cloned from S-specific, class-switched memory B-cells at three visits with median times of 35.5 (Visit 1), 95.5 (Visit 2) and 153.5 (Visit 3) days post-infection. To evaluate functional activity, mAbs were screened at 50 nM (7.5 μg/ml) in a pseudovirus (PV) reporter assay using vesicular stomatitis virus bearing the Wuhan SARS-CoV-2 S (VSV SARS-CoV-2-S). Neutralization was defined as a ≥80% decrease in PV infection relative to virus-only controls, while enhancement was defined as a ≥50% increase in PV infection compared to virus-only controls.

Based on these criteria, neutralizing antibodies comprised 13.0% (158/1213) of the total number of antibodies screened and were found with increasing frequency across Visits 1, 2 and 3 (5.6%, 13.2%, 16.9%, respectively) (**Table 1**). By comparison, infection-enhancing antibodies were found less often, but still comprised 5.9% (72/1213) of all antibodies screened, and increased in frequency from Visit 1 (2.6%) to Visit 3 (7.7%). These findings suggest that SARS-CoV-2 infection-enhancing antibodies are present throughout the COVID-19 convalescent period and persist for at least 5 months post-infection. Further analysis of antibody distribution among donors with different COVID-19 outcomes revealed the presence of infection-enhancing antibodies in individuals with mild, moderate, or severe illness (**Fig. 1A**), suggesting these antibodies are not restricted to those with severe disease and may persist in individuals with a mild course of illness. Infection-enhancing mAbs were identified in all donors at one or more visits (**Fig. 1B**), with higher cumulative numbers across all visits in subjects with severe disease (45/642; 7.0%) as compared to those with mild-to-moderate illness (27/571; 4.7%). However, given the small number of subjects studied, this difference failed to reach statistical significance and it was not possible to determine whether this difference is biologically meaningful.

**Figure 1:**
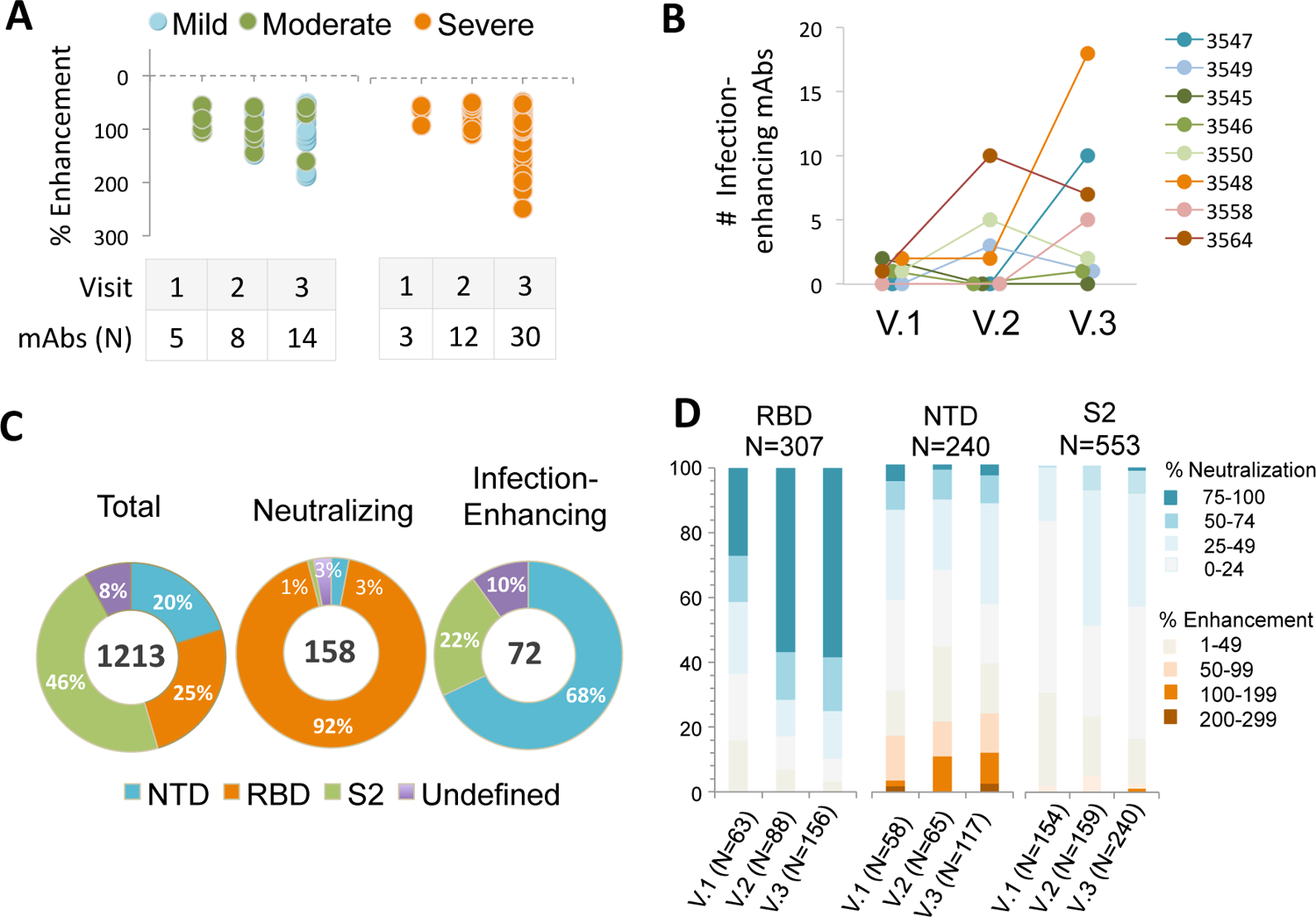
Frequency and binding specificity of SARS CoV-2 infection-enhancing mAbs. **A.** Infection was measured in a VSV-SARS CoV-2-S (Wuhan) pseudovirus (PV) reporter assay in the presence of monoclonal antibodies (mAbs) from eight COVID-19 convalescent subjects with mild (N=2), moderate (N=3) or severe (N=3) illness. Median days post-infection were 35.5 (Visit 1), 95.5 (Visit 2) and 153.5 (Visit 3). Infection enhancement was defined as an increase in infection ≥50% over virus-only controls**. B.** Number of infection-enhancing mAbs identified for individual COVID-19 donors with mild (3547, 3549), moderate (3545, 3546, 3550) or severe (3548, 3558, 3564) illness in longitudinal sampling from Visits 1-3. **C.** Binding specificity of total, neutralizing and infection-enhancing mAbs to recombinant SARS CoV-2 (Wuhan) NTD, RBD or S2 proteins. mAbs were tested from all subjects across all visits. **D.** Proportion of infection-enhancing or neutralizing activities among mAbs binding to recombinant SARS CoV-2 (Wuhan) RBD, NTD or S2. Data represent mAbs from all subjects across visits 1-3.

**Table 1:** Frequency of SARS CoV-2 neutralizing and infection-enhancing mAbs. mAbs cloned a median of 35.5 (Visit 1), 95.5 (Visit 2) and 153.5 (Visit 3) days post-infection from eight COVID-19 convalescent patients.

| SARS CoV-2 Spike (S)-binding mAbs | VISIT 1 | VISIT 2 | VISIT 3 | TOTAL |
| --- | --- | --- | --- | --- |
| Screened (N) | 305 | 334 | 574 | 1213 |
| Neutralizing $\geq 80\%$ (N) (%) | 17 (5.6) | 44 (13.2) | 97 (16.9) | 158 (13.0) |
| Enhancing $\geq 50\%$ (N) (%) | 8 (2.6) | 20 (6.0) | 44 (7.7) | 72 (5.9) |

We next determined the binding specificity of neutralizing and infection-enhancing antibodies to the RBD, NTD and S2 domains. As previously reported (22), the vast majority (92%) of neutralizing mAbs recognized the RBD with a small number binding to regions outside the RBD (**Fig. 1C**). In comparison, the majority of infection-enhancing antibodies were directed to the NTD (68%) with a smaller percentage (21%) recognizing S2. These results are consistent with prior reports describing several FcγR-independent, infection-enhancing mAbs, all of which bound epitopes within the NTD (19,20,23), and further suggest that additional binding specificities associated with infection enhancement may encompass regions of S2.

Further analysis of the functional distribution of neutralizing and infection-enhancing antibodies confirmed that the majority of potent neutralizing activity was associated with mAbs directed to the RBD, while the NTD was targeted by both infection-enhancing antibodies and a small number of neutralizing mAbs (**Fig. 1D**). The S2 domain appeared to be comparatively functionally neutral with neither potent neutralizing nor infection-enhancing mAbs binding to this region, suggesting these antibodies recognize post-fusion epitopes on S2 that are not involved in modulation of virus infectivity.

### Unique VH germline characteristics of NTD-binding and infection-enhancing antibodies

We next determined characteristics of the NTD-binding mAbs by analyzing the variable region of the antibody heavy chain (VH). NTD-binding mAbs were first compared to antibodies directed to the RBD and S2 domains (**Fig. 2**). In agreement with our prior results (22), we found increased representation of mAbs utilizing VH3-30 and VH3-53 germline gene segments among RBD-binding antibodies (**Fig. 2A**). Similarly, S2-binding mAbs utilized a more focused set of VH germline segments including VH1-69, VH3-30 and VH3-30-3. In contrast, NTD-binding mAbs used a diverse array of VH genes, and this broad representation was mirrored in the infection-enhancing antibodies, which utilized VH germline segments including VH1-18, VH1-24, VH1-69, VH2-70, VH3-11, VH3-30, VH3-30-3 and VH4-39. Across all donors, the level of VH somatic hypermutation (SHM) among NTD-binding mAbs increased from Visit 1 to Visit 3 (**Fig. 2B**), with a median number of VH nucleotide substitutions of 2, 6 and 8 for sequential visits. NTD-binding mAbs also displayed increases in binding affinity to a proline stabilized, recombinant SARS-CoV-2 S protein (S2-P) across Visits 1-3 (**Fig. 2C**) consistent with antigen-driven maturation of antibodies to this region.

**Figure 2:**
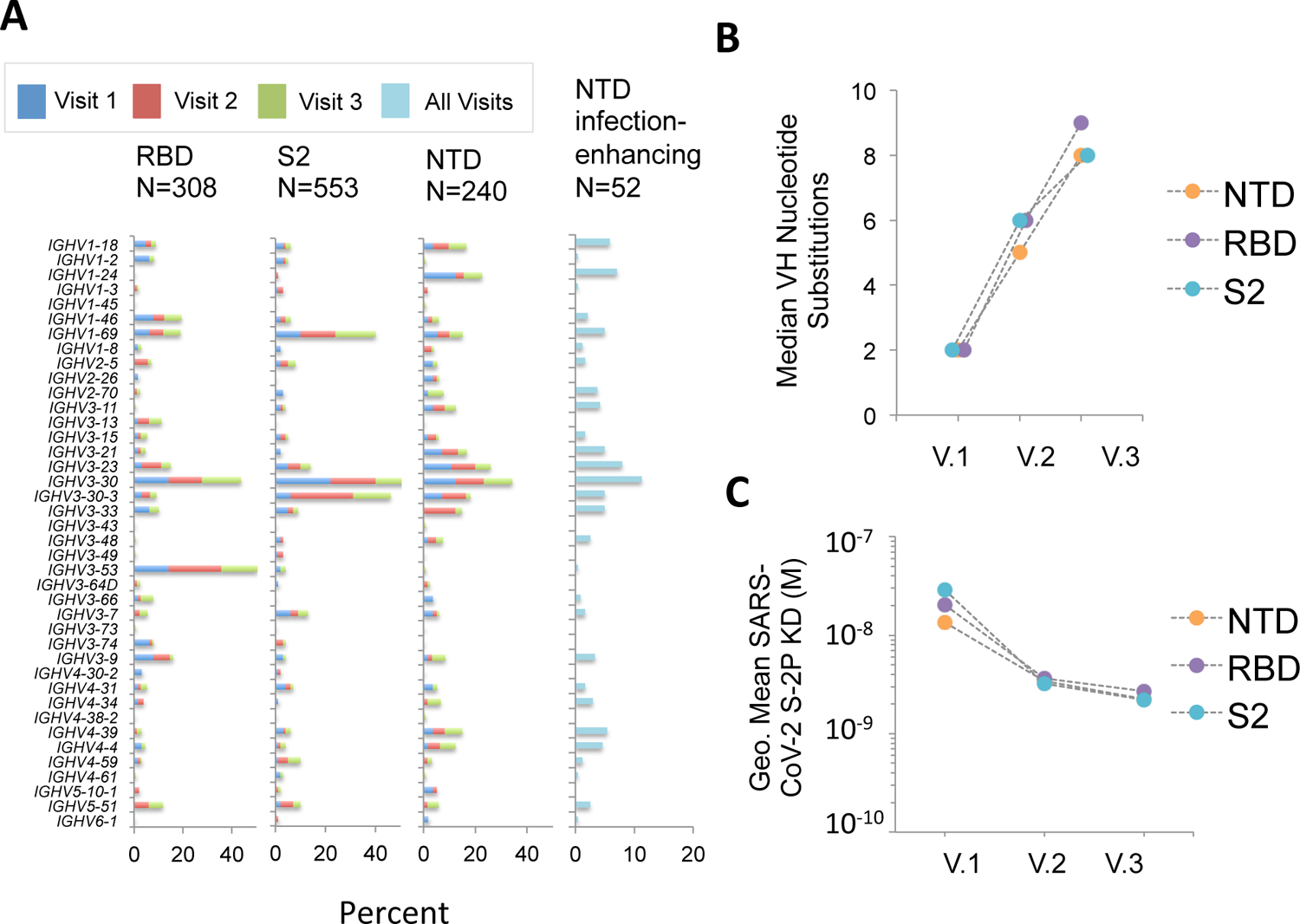
Characteristics of NTD-binding, infection-enhancing mAbs. **A.** Frequency of VH germline usage at Visits 1-3 for mAbs binding to SARS CoV-2 RBD, NTD and S2, and for NTD-binding mAbs with infection-enhancing activity across all visits. **B.** Median number of VH nucleotide substitutions for NTD-, RBD- or S2-specific mAbs from all donors at Visits 1-3. **C.** Geometric mean Fab binding affinities (equilibrium dissociation constant, K_D_) to recombinant, proline-stabilized S (SARS CoV-2 S-2P) for mAbs from all donors at Visits 1-3. MAbs that did not display detectable Fab binding are excluded from this analysis.

### Infection-enhancing activity of NTD-binding mAbs to SARS-CoV-2 variants

We next determined whether the infection-enhancing activity of NTD-binding mAbs also occurred with SARS-CoV-2 VOC. Individual VSV-based PVs were created bearing SARS-CoV-2 S from Beta, Delta, or Omicron (BA.1, BA.2) variants. NTD-binding, infection-enhancing mAbs (N=26) representing clones from multiple donors across Visits 1-3 were tested to determine changes in activity to each variant PV as compared to a Wuhan-PV. The majority of mAbs (24/26) enhanced infection of the Delta-PV at levels comparable to those of the Wuhan-PV, while approximately half (14/26) retained enhancing activity to the Beta-PV (**Fig. 3A**). However, none the mAbs displayed infection-enhancing activity with the BA.1-PV, and only a small number (4/26) enhanced infection of the BA.2-PV. The absence of infection-enhancing activity to the BA.1-PV was associated with near-complete loss of antibody binding to recombinant BA.1 NTD protein (**Fig. 3B**), suggesting that mutations in the NTD of BA.1 abrogate the functional epitopes recognized by these mAbs. Overall, the level of enhancement of infection with PVs was correlated with the level of antibody binding to the respective variant S proteins by ELISA (R^2^=0.546) (**Fig. 3C**), confirming the association of antibody binding with functionality.

**Figure 3:**
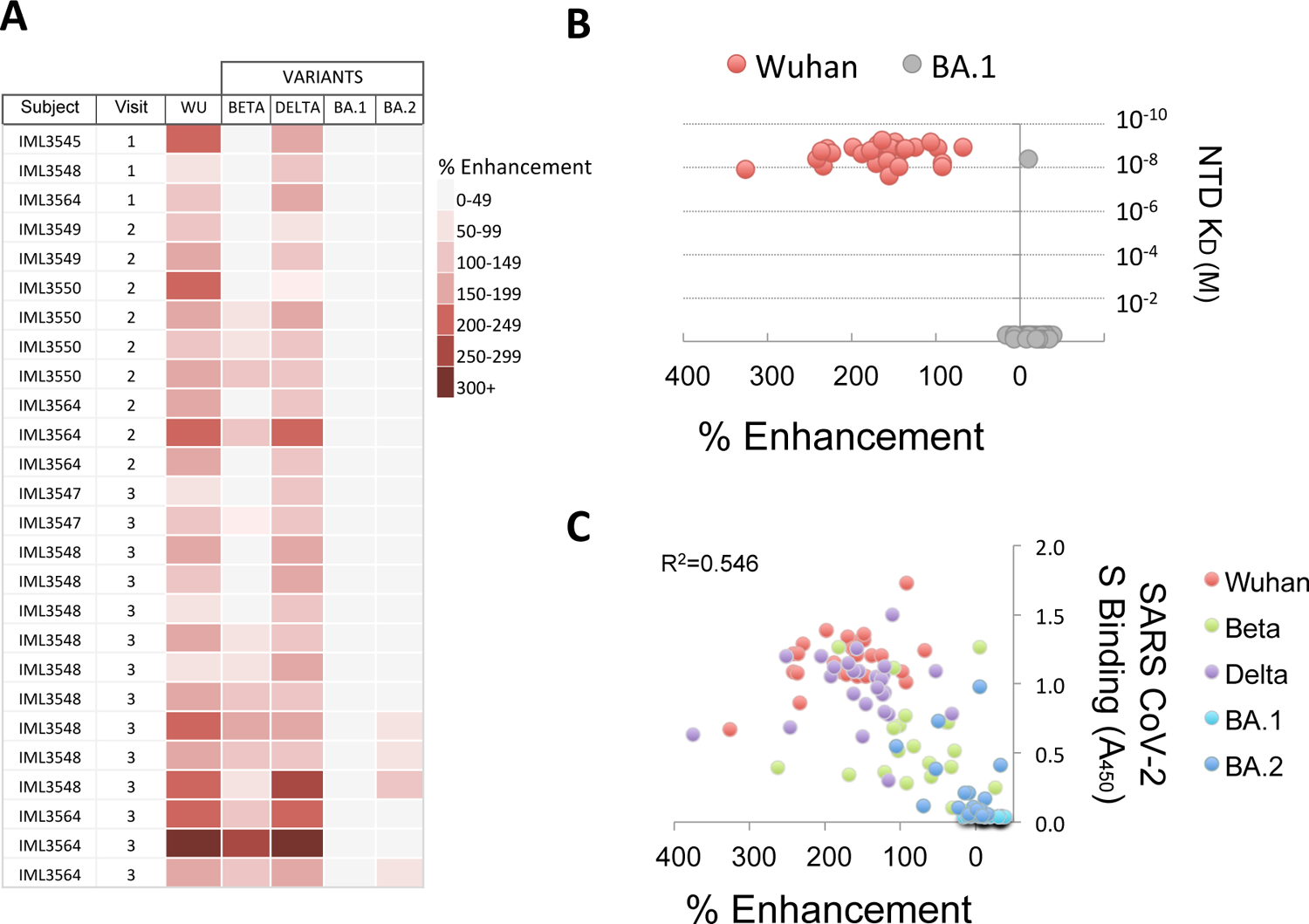
Infection-enhancing activity of NTD-binding mAbs to SARS CoV-2 variants. **A.** NTD-binding mAbs representing clones from multiple donors at Visits 1-3 were tested for infection-enhancing activity in a VSV-SARS CoV-2-S pseudovirus (PV) assay. Infection-enhancement was measured using PV bearing the S from Wuhan (WU), Beta, Delta or Omicron (BA.1, BA.2). **B.** Percent infection enhancement by NTD-binding mAbs to Wuhan- and BA.1-PVs and binding affinities (equilibrium dissociation constant, K_D_) of Fab from each mAb to recombinant Wuhan and BA.1 NTD proteins. **C.** Correlation of percent infection-enhancement and ELISA binding activity (Absorbance 450nm) to recombinant S proteins from Wuhan and SARS CoV-2 variants for NTD-binding mAbs. The coefficient of determination (R^2^) was calculated by linear regression.

### Mechanism of infection-enhancement of SARS-CoV-2

A proposed mechanism for FcγR-independent enhancement of SARS-CoV-2 infection involves bivalent binding of antibodies to the NTD of adjacent S1 subunits, triggering the open conformation of the RBD and increased binding to ACE2 (19). We further probed this mechanism by comparing the enhancing activity of IgG and purified Fab generated from a panel of NTD-binding, infection-enhancing mAbs (**Fig. 4**). Our results demonstrate enhancement of Wuhan-PV infection in the presence of IgG mAbs that was lost when equimolar concentrations of Fab for each mAb were used instead of IgGs, confirming that bivalent antibody binding is required for infection-enhancing activity *in vitro* (**Fig. 4A**). Similar results were found comparing IgG and Fab activities with a Delta-PV, whereas no enhancement was seen with an Omicron BA.1-PV using either IgG or Fab. This latter result is consistent with the observed loss of antibody binding to the BA.1 NTD protein (**Fig. 3B**).

**Figure 4:**
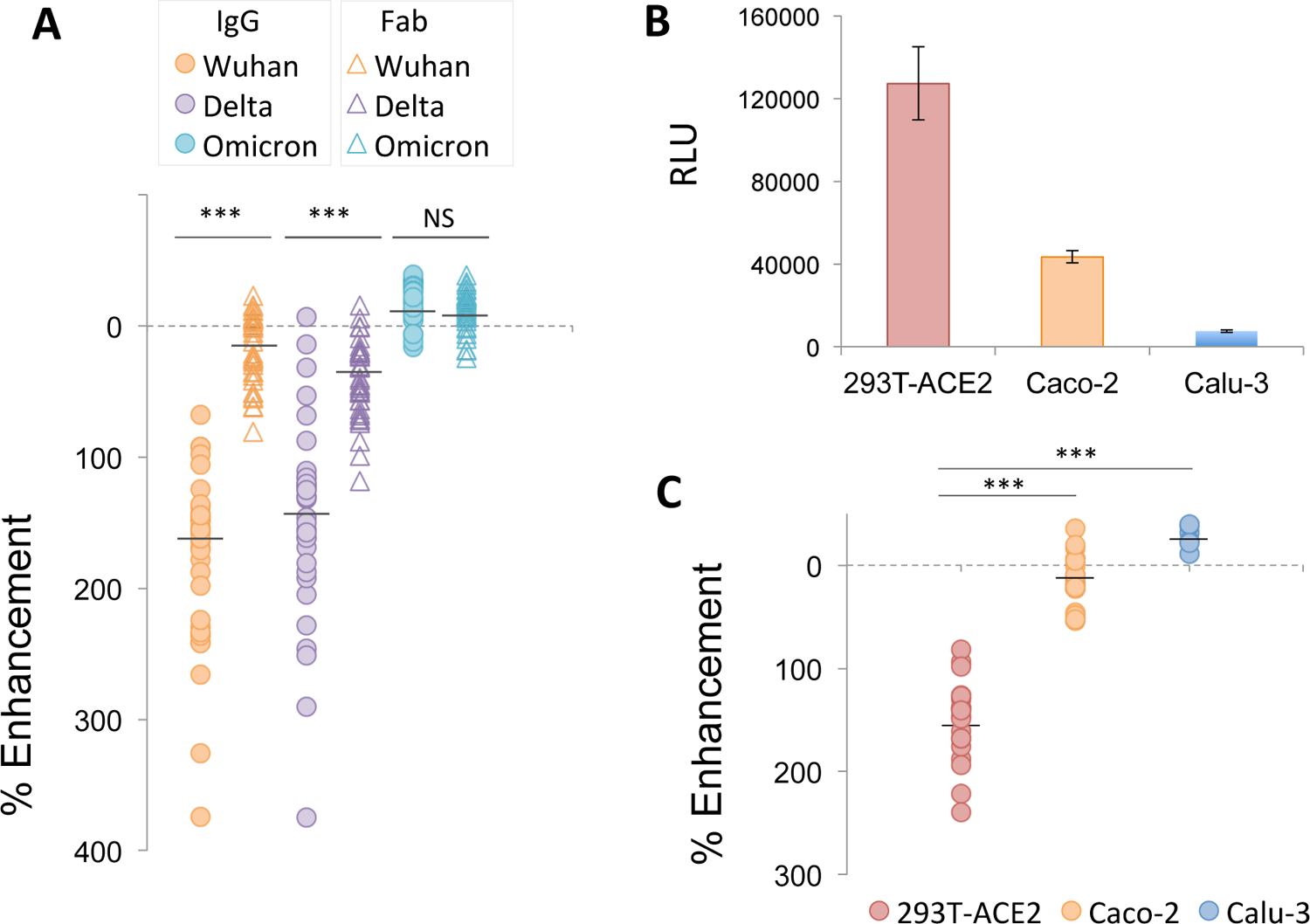
Mechanism of infection-enhancement of SARS CoV-2. **A**. Infection-enhancing activity of IgG and purified Fab generated from a panel of NTD-binding, infection-enhancing mAbs. Enhancement was measured in VSV-SARS CoV-2-S pseudovirus (PV) assays using PV bearing S from Wuhan, Delta or Omicron variants. Mean values for percent enhancement by IgG and Fab against each PV were compared using Student’s t-test, ***p<0.0001, NS=not significant **B.** Infection of 293T-ACE2, respiratory (Calu-3) and intestinal (Caco-2) epithelial cell lines with a Wuhan-PV. Luciferase activity was measured in cell lysates 24 hrs after infection and expressed as the mean ± SD of relative light units (RLU) for triplicate wells. **C.** Percent enhancement of Wuhan-PV infection by NTD-binding mAbs in 293T-ACE2, Calu-3 and Caco-2 cells. Mean values for percent enhancement of infection between two cell types were compared using Student’s t-test, ***p<0.0001.

Our results also confirm that antibody-mediated enhancement of SARS CoV-2 in these assays is independent of engagement with FcR. First, the 293T-ACE2 target cells used for infection do not express FcR and, further, all antibodies used in these experiments were produced in a yeast expression system and are aglycosylated, thus precluding most antibody interaction with FcR. Taken together, these results support a model of bivalent antibody binding to NTD as a necessary component for FcR-independent enhancement of SARS-CoV-2 *in vitro*.

As increased binding to ACE2 is also integral to this proposed mechanism (19), we next evaluated the impact of different ACE2-expressing target cells on the infection-enhancing activity of NTD-binding mAbs. Prior reports of infection-enhancing mAbs (19), as well as our own assays (22,24,25), utilize 293T cells that overexpress ACE2 as targets for SARS-CoV-2 PV infection. In comparison, biologically relevant target cells for SARS-CoV-2 include epithelial cells lining the respiratory and gastrointestinal tracts, which may express lower levels of ACE2 (26–29). We found respiratory (Calu-3) and intestinal (Caco-2) epithelial cell lines yielded significantly lower levels of PV infection as compared to 293T-ACE2 cells after 24 hrs *in vitro* (**Fig. 4B**). Moreover, when compared with virus-only controls in each cell type, we found no evidence of infection enhancement by NTD-binding mAbs in either Calu-3 or Caco-2 cells (**Fig. 4C**). This finding suggests that antibody-mediated enhancement of SARS-CoV-2 may be conditional upon the choice of target cells, and may be less likely to occur in epithelial cells associated with the respiratory or gastrointestinal tracts.

### Expanded functional activity of NTD-binding serum mAbs

The significance of antibody-mediated enhancement to the pathogenesis of SARS-CoV-2 has been called into question following results in animal models, which found no evidence of enhanced disease in the majority of animals infused with an NTD-binding, infection-enhancing mAb prior to virus challenge (20). However, other studies in humans have demonstrated increased frequency of these antibodies in the sera of COVID-19 patients with severe disease (19). We sought to examine the functionality of serum antibodies from two of the eight COVID-19 convalescent patients in our cohort. These two patients (3546, 3548) had moderate and severe illness, respectively, and were evaluated within the context of a study of their serum antibody repertoires to SARS-CoV-2 (30). A total of twelve unique mAbs, representing the most abundant serum clonotypes in each subject, were recombinantly expressed in Expi293 cells (30). These antibodies are fully glycosylated and capable of engagement with FcγR. Five were found to bind the NTD, of which four enhanced infection with SARS-CoV-2 PVs *in vitro* (30) (**Fig. 5A**).

**Figure 5:**
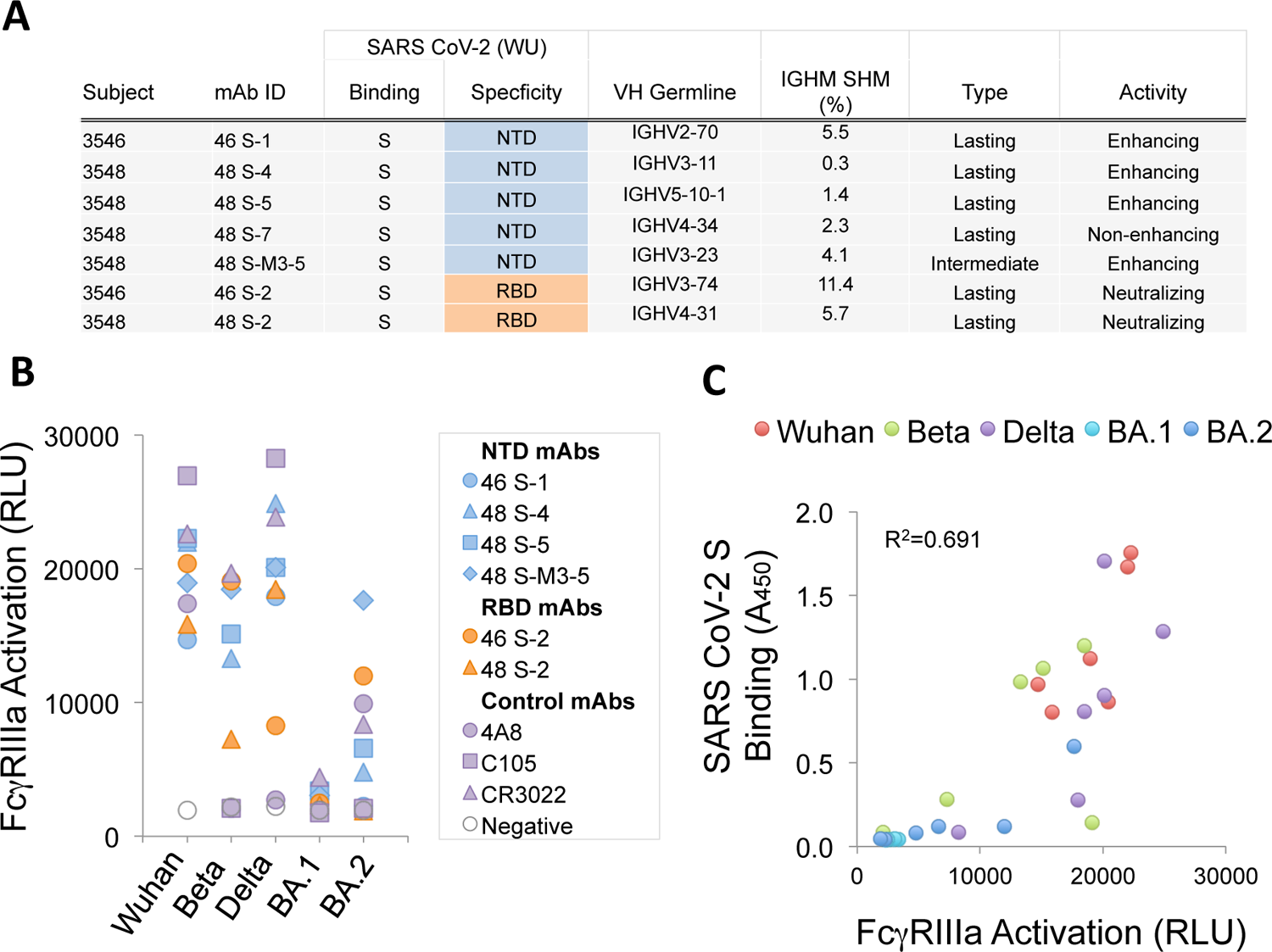
Features and functional activity of NTD-binding, infection-enhancing serum mAbs. **A.** IgG mAbs representing major serum clonotypes from two donors (3546, 3548) were characterized for SARS CoV-2 binding specificity, variable heavy chain (VH) germline use, percent VH somatic hypermutation (IGHM SHM) and categorized as lasting (detected at Visit 3), intermediate (detected at 2, but not Visit 3), or transient (detected only at Visit 1) as described (30). Functional activity was determined using a VSV-SARS CoV-2-S (Wuhan) pseudovirus (PV) assay to measure neutralization or infection-enhancement as reported (30). **B.** Serum mAbs binding NTD (46 S-1, 48 S-4, 48 S-5, 48-S-M3-5), RBD (46 S-2, 48 S-2), or control mAbs (4A8, C105, CR3022) were tested for activation of FcγRIIIa in a luciferase reporter-cell assay with recombinant S from Wuhan, Beta, Delta or Omicron (BA.1, BA.2). FcγRIIIa activation was measured as relative light units (RLU). **C.** Correlation of FcγRIIIa activation (RLU) and SARS CoV-2 S binding by ELISA (Absorbance 450 nm) for NTD- and RBD-binding mAbs to S from Wuhan, Beta, Delta or Omicron (BA.1, BA.2). The coefficient of determination (R^2^) was calculated by linear regression.

The four NTD-binding, infection-enhancing mAbs were evaluated for additional functionality by measuring their ability to activate FcγRIIIa expressed on reporter cells as a surrogate measure of antibody-dependent cellular cytotoxicity (ADCC). Control antibodies included 4A8, an NTD-specific, neutralizing mAb (13), two RBD-specific, neutralizing mAbs from the same subjects (46 S-2, 48 S-2), and two additional RBD-specific neutralizing mAbs, C105 and CR3022 (31–33). In assays using recombinant S proteins as antigens, all four serum-derived NTD-binding mAbs activated FcγRIIIa at levels comparable to serum-derived RBD-binding mAbs and to the positive control antibodies (**Fig. 5B**). Evaluation of additional S proteins from Beta, Delta, and Omicron BA.1 and BA.2 variants revealed diminished responses by some mAbs to Beta-S and complete loss of FcγRIIIa activation to BA.1-S by all mAbs. FcγRIIIa activation was correlated with the level of antibody binding to the recombinant S proteins from each variant by ELISA (R^2^=0.691) (**Fig. 5C**), suggesting that mutations in epitopes recognized by these mAbs are responsible for the loss of functionality. These results are consistent with a prior study of FcR-mediated effector function by an NTD-binding mAb (21) and further suggest that NTD-binding antibodies that display infection-enhancing properties *in vitro* also have the capacity to activate FcγRIIIa and may therefore have the potential to mediate ADCC. These results also align with data demonstrating additional FcγR-mediated effector functions associated with these same serum-derived mAbs, including antibody-dependent cellular phagocytosis (ADCP) and antibody-dependent complement deposition (ADCD) (30).

## DISCUSSION

Characterization of the functional antibody responses to the SARS-CoV-2 S protein is important for understanding the immune correlates of protection and for informing vaccine development. Most current vaccine platforms focus on induction of neutralizing antibodies to the RBD of the S1 subunit, while less attention has been paid to the characteristics and functional attributes of antibodies to the S1 NTD. Both neutralizing (13–18) and infection-enhancing antibodies (19,20,23) directed to the NTD have been described, but their role, if any, in modulating SARS-CoV-2 infection and disease is less clear. In the present study, we examined antibody responses to the NTD using mAbs derived from longitudinal sampling of B-cells collected from eight COVID-19 convalescent patients. Our results reveal several novel aspects of the antibody response to the NTD and the functionality of NTD-binding mAbs.

First, by screening a large collection of S-specific mAbs derived from the memory B-cell populations of donors with mild, moderate, or severe COVID-19, we found infection-enhancing mAbs present in all donors irrespective of disease severity, and these antibodies persisted for up to five months post-infection. Infection-enhancing mAbs represented 5.9% of the total number of antibodies screened—roughly half the number of neutralizing mAbs identified— suggesting they may constitute a more significant pool of antibodies than previously recognized. Moreoever, as compared to the RBD and S2 domains, we found broader representation of VH germline segments among antibodies binding the NTD, suggesting the possibility of greater diversity among NTD-binding specificities. Functional analysis further supported this observation by revealing both neutralizing and infection-enhancing activities associated with NTD-binding mAbs as compared to RBD-binding mAbs, which represented the majority of neutralizing activity. Overall, these findings suggest the NTD elicits antibodies with the potential for diverse functionality *in vivo*.

Next, we confirmed and extended results on the proposed mechanism of FcγR-independent enhancement of SARS-CoV-2 infection (19). Our findings support observations that infection enhancement is mediated predominantly by NTD-binding mAbs and is predicated on bivalent antibody binding to S. Antibody binding is modeled to trigger the open conformation of RBD and to engage multiple ACE2 receptors on the host cell surface (19). Earlier studies demonstrated enhancement of SARS-CoV-2 infection by NTD-binding mAbs in high-ACE2 expressing 293T cells and Huh7 cells, a human liver carcinoma cell line that expresses ACE2 (19,34). However, when using epithelial cell lines derived from human lung (Calu-3) and intestine (Caco-2), we found no evidence of SARS-CoV-2 infection enhancement by NTD-binding mAbs. Both Caco-2 and Calu-3 cells express ACE2 and are widely used to characterize SARS-CoV-2 infection *in vitro* (35–39). The relative absence of infection enhancement with NTD-binding antibodies in these cells may result from lower ACE2 expression, a possibility that aligns with the observed impact of ACE2 expression levels on experimental measures of antibody neutralization of SARS-CoV-2 (40). When considered alongside the functional requirement of bivalent antibody binding to NTD, it is possible that limiting levels of ACE2 on target cells may preclude active engagement of multiple receptors, thereby abrogating any enhancing effect of these antibodies on virus infectivity. While we cannot rule out post-entry effects on PV infection in these cells, our findings suggest the choice of target cell plays an important role in the phenotypic expression of FcγR-independent antibody enhancement of SARS-CoV-2, and raise the possiblity that enhancement may be less likely to occur in epithelial cells in the respiratory or gastrointestinal tracts. Overall, it remains unclear whether this mechanism is widely, or even functionally, operative *in vivo*, particularly in light of results from animal studies, which found limited evidence of SARS-CoV-2 antibody-mediated disease enhancement (20).

Our results also demonstrate loss of NTD antibody binding and functional activity to SARS-CoV-2 Omicron BA.1 and BA.2 variants. Comparison of NTD mutations in BA.1 relative to ancestral SARS-CoV-2 demonstrates multiple changes in positions 211-214, which encompass residues reported to impact binding of infection-enhancing mAbs (**Fig. 6**) (19,20). Mutations in this region are also present in the NTD of Beta (D215G) and Omicron BA.2 (V213G), of which the latter change is maintained in all recent Omicron lineages, including BA2.75, BA 4/5, BQ1.1 and XBB (**Fig. 6A**). Codons from 213-216 are also sites of recurring insertion elements among different SARS-CoV-2 lineages (41); however, it is not known whether these recurrent insertions arise in response to immune selection or reflect compensatory changes that favor increased virus infectivity (41). Additional mutations in the NTD spanning positions 24-27 also persist in all Omicron lineages and these are directly adjacent to residues 27-32 implicated in the binding of infection-enhancing mAbs (20). Taken together, mutations in the NTD constituting the neutralization supersite (residues 14–20, 140–158 and 245–264) and those implicated in binding infection-enhancing mAbs (19,20) constitute most of the persisting changes in the NTD sequence of SARS-CoV-2 variants (**Fig. 6A**) and impact discrete domains in the NTD protein (**Fig. 6B**). While counterintuitive, our results support the thesis raised by others (19) that certain NTD-binding mAbs, while mediating enhanced infection with SARS-CoV-2 *in vitro*, may have protective FcγR-effector functions *in vivo*, with the potential to drive immune selection and NTD sequence variability at this site.

**Figure 6:**
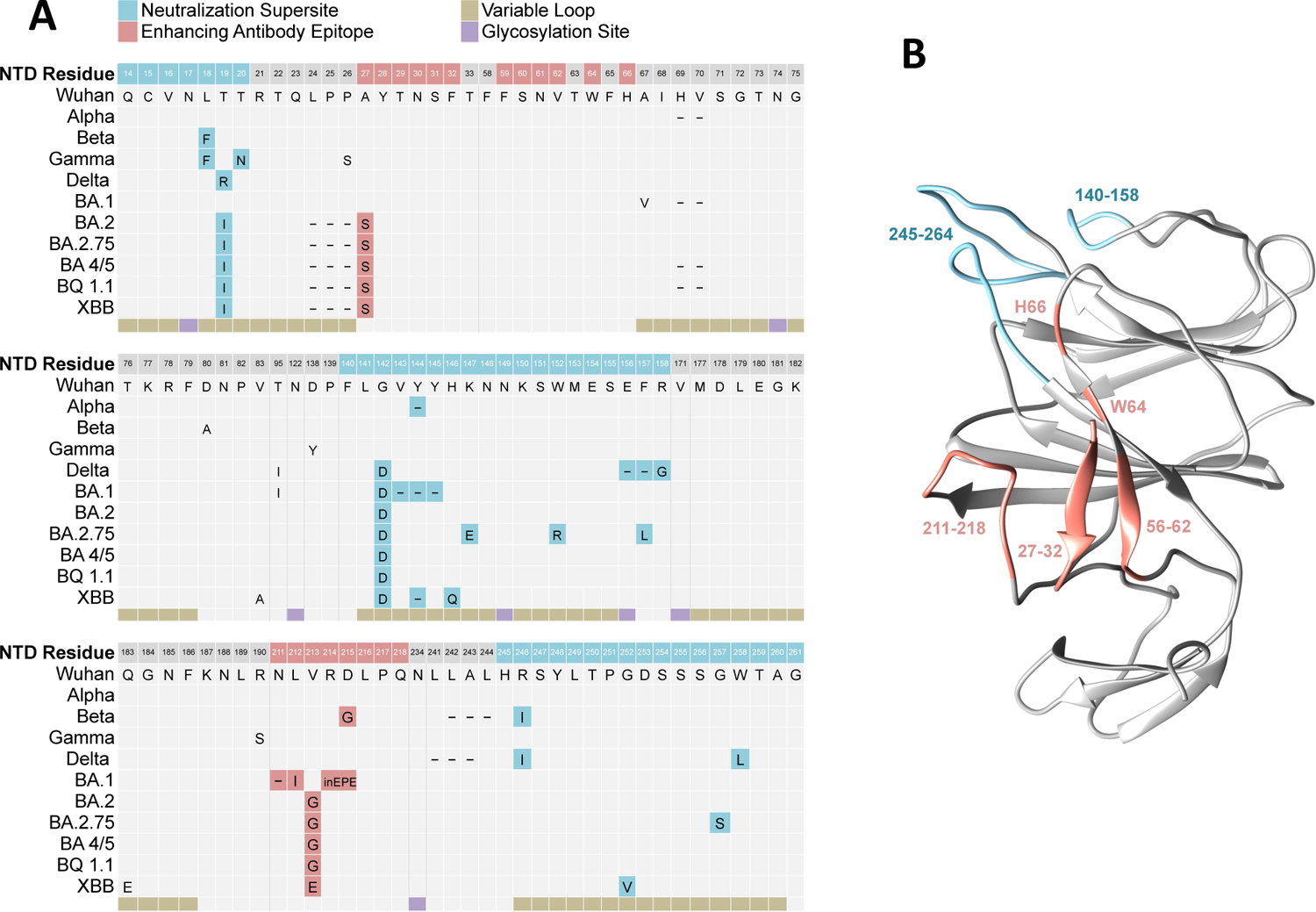
Mutations in the S1 NTD of SARS CoV-2 variants of concern. **A.** Sequence alignment of the N-terminal domain (NTD) of the SARS CoV-2 spike protein from ancestral (Wuhan) and variant strains. Structural features include five hypervariable loops (beige) and glycosylation sites (purple). Mutations mapping to regions within the convergent neutralization supersite (blue) and residues implicating in the binding of infection-enhancing mAbs (red). **B.** Model of the three-dimensional structure of the SARS CoV-2 NTD highlighting residues involved in antibody-mediated neutralization (blue) or infection-enhancement (red). Visualization was performed using UCSF Chimera and unresolved loops were modeled using the Modeller web server (47,48).

Importantly, our results demonstrate for the first time that NTD-binding, infection enhancing antibodies are present as major serum clonotypes that can last in circulation for months during COVID-19 convalescence. Prior reports have described infection-enhancing mAbs derived from SARS CoV-2 S-reactive plasmablasts or memory B cell populations (19,20); ours is the first to identify NTD-binding, infection-enhancing mAbs isolated directly from serum (*i.e*., functionally secreted antibody). While these serum-derived NTD-binding antibodies enhance SARS-CoV-2 PV infection *in vitro*, they also mediate activation of FcγRIIIa in response to binding SARS-CoV-2 S proteins, suggesting the potential for broader functionality that may include protective ADCC responses *in vivo*. FcγRIIIa activation by these antibodies was almost completely abrogated to BA.1 and BA.2 S proteins, again suggesting that mutations in the regions of the NTD recognized by these antibodies may contribute to a loss of functionality.

Overall, our results suggest that enhancement of SARS-CoV-2 infection in the context of NTD-binding mAbs may be a unique phenomenon observed under certain *in vitro* conditions. This attribute, along with reports of their characteristic downward angle of binding to the SARS-CoV-2 S protein (19,20) may help distinguish these mAbs from other NTD-binding antibodies, including those with neutralizing activity. In a broader context, our data support the concept that these antibodies arise during natural infection and represent a significant proportion of the memory B-cell antibody repertiore during convalescence. Evidence from this study demonstrates these NTD-binding, infection-enhancing mAbs can also be present as major secreted clonotypes in serum, they are able to activate cells through FcγRIIIa, and raise the possibility they may mediate protective FcR-effector functions *in vivo*. Further studies are needed to determine whether these antibodies contribute to immune protection, and whether they are induced by natural infection with SARS-CoV-2 variants or by immunization with next generation multivalent vaccines.

## MATERIALS AND METHODS

### Study participants and design

Eight subjects with PCR-confirmed SARS-CoV-2 infection were recruited for this study from 8 April to 22 May, 2020 (24). Based on their infection dates, it is presumed the infecting strain was a Wuhan-Hu-1 (Wuhan) isolate. The clinical and biological characteristics of the participants have been reported in detail elsewhere (22,24,25,42). As per the study design, participants were enrolled within approximately 1 month (median 35.5 days) after SARS-CoV-2 diagnosis and were followed for at least 6 months. Two additional specimen collection visits occurred at approximately 2 months (Visit 2, median 95.5 days) and 5 months (Visit 3, median 153.5 days) after the initial visit. Based on clinical presentation and medical history, five patients were considered to have mild to moderate COVID-19 illness, and three had severe illness with hospitalization. Blood samples were processed to isolate B cells and serum in the Immune Monitoring Core Laboratory at the Dartmouth Geisel School of Medicine. The isolated B cells and serum were aliquoted and stored at −80°C until further use. The study was approved by the Dartmouth-Hitchcock Hospital (D-HH) Human Research Protection Program (Institutional Review Board) and all volunteers gave informed consent using an approved D-HH template.

### Expression and purification of IgGs and Fab fragments

Monoclonal antibodies used for binding and functional assays were produced as full-length IgG1 proteins in *S. cerevisiae* cultures, as previously described (1). Briefly, yeast cultures were incubated in 24-well plates at 30°C and 80% relative humidity with shaking at 650 RPM in Infors Multitron shakers. After 6 days of growth, the culture supernatants were harvested by centrifugation and IgGs were purified by protein A-affinity chromatography. IgGs bound to the agarose were eluted with 200 mM acetic acid with 50 mM NaCl (pH 3.5) and neutralized with 1/8 (v/v) 2 M HEPES (pH 8.0).

To generate Fab fragments, IgGs were digested with papain for 2 hrs at 30°C followed by the addition of iodoacetamide to terminate the reaction. The mixtures were then passed over protein A agarose to remove Fc fragments and undigested IgG. The flow-through of the protein A resin was passed over either KappaSelect resin or LambdaFabSelect resin (Cytiva) for antibodies utilizing the kappa or lambda light chain classes. The Fabs captured on the resin surface were eluted using 200 mM acetic acid/50 mM NaCl (pH 3.5) into 1/8th volume 2 M Hepes (pH 8.0). Monovalent equilibrium dissociation constant (K_D_) affinities of Fabs were measured using biotinylated antigens (100 nM, 1.0-2.0 nm) immobilized on streptavidin biosensors (Molecular Devices) as described (22).

### Isolation and characterization of serum mAbs

Methods for the isolation and characterization of serum antibodies to SARS CoV-2 from these patients are described in detail in a companion paper (30). In brief, serum IgG was enriched by passing serum through Protein G agarose columns and eluting the bound antibodies with 100 mM glycine-HCl, pH 2.7. Purified IgG was digested into F(ab’)2 with 25 µg of IdeS protease per 1 mg of IgG and then incubated with Strep-Tactin agarose (IBA-Lifesciences) for 1 hr to remove the IdeS protease. For each sample, F(ab’)2 were further enriched by passage through affinity columns of N-hydroxysuccinimide (NHS)– activated agarose resin coupled with recombinant Wuhan SARS CoV-2 S or RBD proteins as described (30). Samples were then denatured/alkalized, followed by digestion with trypsin (1:30 (w/w) trypsin/protein) for 16 hrs at 37°C. The resulting peptides were purified using a Hypersep SpinTip C-18 (Thermo Fisher Scientific) and analyzed by liquid chromatography-tandem mass spectrometry (LC-MS/MS). Sequence data for V_H_ and V_L_ from each donor was used to construct protein sequence databases. V_H_ sequences were then grouped into clonotypes based on hierarchical clustering. Selection of antibody sequences for recombinant expression was based on the combination of V_H_:V_L_-paired databases and proteomics data. These genes were then purchased as eBlocks (Integrated DNA Technologies) and cloned into the pcDNA3.4 vector (Invitrogen). Heavy and light chain plasmids for each mAb were transfected into Expi293 cells, and the expressed antibodies purified from cell culture supernatants using Protein G agarose. Using a matched donor-specific database of putative B cell receptor sequences at Visit 1, the secreted NTD-binding mAbs were matched to mAbs cloned from the memory B-cell population of each subject (30). Recombinant antibodies representing each of the 12 most abundant serum IgG clonotypes from two donors with moderate and severe symptoms (Subjects 3546 and 3548, respectively) were characterized by ELISA and biolayer interferometry (BLI) for binding specificity, and for neutralization and Fc-mediated functional activities as described (30).

### VSV-SARS-CoV-2 pseudovirus neutralization assay

Neutralization assays were performed using a VSV-SARS-CoV-2 pseudovirus system as previously described (22,24,25,43). Plasmids expressing the SARS-CoV-2 S protein were obtained from Addgene: pcDNA3.3-SARS2-B.1.617.2 (Delta) was a gift from David Nemazee (Addgene plasmid #172320; http://n2t.net/addgene:172320; RRID:Addgene_172320) (44); pTwist-SARS-CoV-2D18 B.1.1.529 (Omicron) was a gift from Alejandro Balazs (Addgene plasmid #179907; http://n2t.net/addgene: 179907; RRID:Addgene_179907 (45); pcDNA3.3_SARS2_omicron BA.1 (Addgene plasmid #180375; http://n2t.net/addgene: 180375; RRID:Addgene_180375) and pcDNA3.3_SARS2_omicron BA.2 (Addgene plasmid #183700; http://n2t.net/addgene: 180700; RRID:Addgene_180700) were gifts from David Nemazee (46) and were used to create VSV-SARS-CoV-2-S pseudoviruses (43). The pcDNA3.3 SARS CoV-2-Beta S expression plasmid was created in-house by site-directed mutagenesis of pcDNA3.3 SARS CoV-2 S to incorporate mutations (L18F, D80A, D215G, Δ242-244, R246I, K417N, E484K, N501Y/T, D614G and A701V) associated with the Beta variant S. To measure neutralization, individual mAbs were diluted to 50 nM (7.5 μg/ml) and incubated with VSV-SARS-CoV-2-S pseudoviruses for 1 hr at 37°C before adding to either 293T-hsACE2 (Integral Molecular, Philadelphia, PA), Caco-2 (ATCC HTB-37) or Calu-3 (ATCC HTB-55) cells The cells were incubated at 37°C, 5% CO_2_ for 24 hrs, after which luciferase activity (Relative Light Units, RLU) was measured in cell lysates using the Bright-Glo system (Promega, Madison, WI) with a Bio-Tek II plate reader. Percent neutralization was calculated as 100-(mean RLU test wells/mean RLU positive control wells) x 100.

### Monoclonal antibody binding ELISA

For antibody binding studies, 96-well ELISA plates were coated with 50 μl per well of recombinant SARS-CoV-2 (Wuhan) NTD, RBD, S2 proteins, or SARS-CoV-2 S proteins from Wuhan, Beta, Delta, Omicron BA.1 or Omicron BA.2 (Immune Technology) diluted to 5 μg/ml in 1X PBS and incubated overnight at 4 °C. Wells were washed 3 times with PBS and blocked with 5% non-fat dry milk (NFDM)-PBS for 1 h at 37 °C. After removal of blocking buffer, test mAbs diluted to 100 nM in 5% NFDM-PBS were added and incubated for 1 h at 37 °C. Plates were then washed three times with PBS and secondary cross-adsorbed anti-human Fab-HRP (Jackson ImmunoResearch Laboratories) detection antibody was added at 1:10000 dilution in 5% NFDM-PBS for 1 hr at 37 °C. After washing three times with PBS, 50 μl per well of the 1-Step™ Ultra TMB-ELISA Substrate Solution (Thermo Fisher Scientific) was added to detect binding followed by addition of an equal volume of the stop reagent as per manufacturer recommendations. Absorbance was measured at 450 nm wavelength using a Bio-Tek II plate reader.

### FcγRIIIa activation reporter assay

The ADCC potential of individual monoclonal antibodies was measured as described (25) using a Jurkat Lucia NFAT cell line (Invivogen, jktl-nfat-cd16), cultured according to the manufacturer’s recommendations, in which engagement of FcγRIIIa (CD16) on the cell surface leads to the secretion of luciferase. One day prior to running the assay, a high binding 96-well plate was coated with 1 µg/mL SARS-CoV-2 S at 4°C overnight. Plates were then washed with PBS + 0.1% Tween20 and blocked at room temperature for 1 hr with PBS + 2.5% BSA. After washing, dilute serum or nasal wash sample and 100,000 cells/well in growth medium lacking antibiotics were cultured at 37°C for 24 hrs in a 200 µl volume. The following day, 25 µL of supernatant was drawn from each well and transferred to an opaque, white 96 well plate, to which 50 µL of QuantiLuc substrate was added and luminescence immediately read on Bio-Tek II plate reader. Negative control wells substituted assay medium for sample while 1x cell stimulation cocktail (Thermo Fisher Scientific) plus an additional 2 µg/mL ionomycin were used to induce expression of the transgene as a positive control.

## ACKNOWLEDGEMENTS

The authors acknowledge the Immune Monitoring and Flow Cytometry Resource (IMFCSR) at the Norris Cotton Cancer Center at Dartmouth supported by NCI Cancer Center Support Grant 5P30 CA023108-41. The research was supported by the Bill and Melinda Gates Foundation (Grant Number DGR5180); Dartmouth-Hitchcock Department of Medicine and Division of Infectious Disease and International Medicine (SEAM Award AY21); and NIH (Grant Number P20 GM113132 to J.L.). VSV-g and SARS-CoV-2-S expression plasmids were generously provided by Dr. Michael Letko (Rocky Mountain Laboratories). The authors thank Alejandra Prevost-Reilly, Dr. Anais Ovalle, and Dr. David de Gijsel for support with clinical specimen collection and donor enrollment.

